# MTA influences RNA Polymerase II transcription dynamics and regulates the cold response in Arabidopsis

**DOI:** 10.1101/2023.05.10.540235

**Authors:** Susheel Sagar Bhat, Dawid Bielewicz, Peter Kindgren

**Affiliations:** Umea Plant Science Centre, Department of Forest Genetics and Plant Physiology, Swedish University of Agricultural Sciences, 90187 Umea, Sweden; Centre of Advanced Technology, Adam Mickiewicz University Poznan, 61-614 Poznan, Poland

## Abstract

RNA methylation at the N6 position of adenosine residues (m^6^A) has been demonstrated to be a vital epigenetic mark in eukaryotes. In plants, it is involved in a myriad of developmental and stress responsive pathways, such as cold stress. However, the role of MTA (m^6^A writer) itself in such processes remains obscured. Here, we show that the cold induced changes in abundance of differentially methylated transcripts are at-least partly due to the differences in the rates of transcription of such transcripts. RNAPII stalling was observed only on RNA methylation motifs that were decorated with m^6^A, and those that had a specific nucleotide composition. We note that the downregulation of genes in a *mta* mutant is a result of lower transcription of these genes. Moreover, 5’ ends of transcripts from a subset of genes are disproportionately more accumulated in *mta* as compared to wild type under cold stress. In the *mta* mutant, genes corresponding to these transcripts had significantly reduced RNAPII occupancy on their 3’ ends but not on 5’ ends indicating that *mta* serves as an important factor for RNAPII elongation in a subset of genes. Taken together, our data suggest a novel direct role for MTA as a gene specific influencer of RNAPII elongation dynamics and thus introduces a new path by which the m^6^A methylation machinery may affect differential gene expression during stress in plants.

## INTRODUCTION

m^6^A is the most prevalent epigenetic mark found in eukaryotic mRNAs (1). mRNA adenosine methylase (MTA) along with methyltransferase B (MTB), FKBP12 interacting protein 37kDa, Virilizer (VIR) and HAKAI form the m^6^A methyltransferase complex that deposits m^6^A on mRNAs (2–4). MTA is known to be the catalytic protein of the ‘writer’ complex, however, recently MTB has also been shown to have *in vitro* methyltransferase activity (5). Virtually all proteins in the methyltransferase complex are conserved among mammalian and plant systems. Mammalian homologs of the complex are also critical for sustenance of life and disrupting the complex is lethal [reviewed in (6)]. m^6^A can also be found in non-coding RNAs (ncRNAs) like primary-microRNAs (7–9) and novel proteins with m^6^A methyltransferase activity have been identified including methyltransferase like 16 (METTL16) and METTL5 [reviewed in (10)]. Plant primary-microRNAs are also known to be methylated by MTA (11). Apart from the ‘writer’ complex, ‘reader’ and ‘eraser’ proteins are implicated in the downstream pathways that eventually lead to observable m^6^A related effects. A specific sequence motif, RR**A**CH (R = A/G, H = A/C/U and **A** being the methylated adenosine), is known to be enriched around the m^6^A peaks with UGUA being one of the most abundant motifs in plants while GGAC/GAAC motifs are known to be more prevalent in mammalian systems (8, 9, 12–14). Notably, not all RRACH motifs carry the m^6^A mark which is known to be reversible (‘erasers’ possess demethylase activity). The mechanisms behind this differential methylation of certain RRACH motifs are still unclear. A mammalian study found that METTL14 (homolog of MTB) guides the deposition of m^6^A by binding to the histone H3 trimethylation at Lys36 (H3K36me3) mark in the chromatin (15). Similarly, gene structures like exon-intron junctions and interactions with other transcription factors may affect the selection of m^6^A motif to be methylated (16). One may speculate that demethylation of certain motifs may contribute to the motif specificity, however, a mammalian study provided evidence that shows that m^6^A content of newly synthesized RNA (chromatin associated RNA) and mature mRNA is almost identical (∼90% overlap), indicating that once deposited m^6^A marks are sustained for majority of a transcript’s life (17).

In plants, changes in m^6^A methylation levels have been shown to affect development and response to stress. Null mutations of all known members of the ‘writer’ complex, with the exception of HAKAI, are embryo lethal pointing towards their critical role in plant survival. Mutants with decreased expression of m^6^A ‘writer’ complex proteins have been produced and show abnormalities in various phenotypic traits like trichome morphology, meristem maintenance, flowering time etc. (2–5, 18). Similarly, m^6^A is of critical importance for proper stress response. Under salt stress, transcripts required for proper salt-stress response are stabilized by deposition of m^6^A marks while the negative regulators of salt stress are stabilized by the loss of m^6^A (13, 19). Furthermore, an increase in m^6^A leads to a decrease in the prevalence of RNA secondary structure which eventually alters the binding of proteins and protein production from such targets (20). A m^6^A demethylase AlkB homolog 9B (ALKBH9B) has been shown to be involved in plant response to viral infection. Interestingly, ALKBH9B does not target endogenous Arabidopsis RNA for demethylation, only viral RNA is targeted. An increase in the m^6^A methylation of the viral RNA makes it more susceptible to non-sense mediated decay (21). Recently, m^6^A was shown to be involved in cold stress response in plants. *mta* hypomorphic mutant is hypersensitive to cold and changes in the m^6^A landscape were observed upon cold stress. Notably, m^6^A peaks around 5’ ends and in the CDS region showed more variation in cold than those located at the 3’ ends. Transcripts that gained m^6^A methylation under cold stress showed increased abundance and polysome association (12).

Aberrant temperature changes pose a significant challenge to plants and are usually dealt with an array of stress responses usually initiated by physiological and transcriptional changes. Cold stress, experienced by plants exposed to low (but not freezing) temperatures results in the activation of *cold responsive (COR)* genes in either a C-repeat binding factor (CBF) dependent or independent pathway (22). As key transcription factors, CBF1, CBF2 and CBF3 have been shown to be induced as soon as 15 min after exposure to cold (23–25). The *CBF* genes sit in tandem on chromosome 4 in *Arabidopsis* and have been shown to act redundantly with at least 65% of *COR* genes regulated by two or three CBFs (26). Nevertheless, the *CBF* region is the most induced genomic region in Arabidopsis in response to cold temperatures (27) and crucial for the cold acclimation process (23, 25).

In this study, we provide evidence that MTA protein itself and not just the m^6^A mark plays an important role in the cold acclimation process in Arabidopsis. We use RNA-seq and plant Native Elongation Transcript sequencing (plaNET-seq) data to show that cold induced changes in the methylation statuses of transcripts are correlated to their transcription levels. We provide evidence that RNA polymerase II (RNAPII) stalling over any given methylation motif is selective and depends on the initial nucleotides in any given motif. The stalling pattern of RNAPII around the methylation motif and the accumulation of 5’ ends of transcripts in the *mta* mutant under cold stress points to the role of MTA in influencing RNAPII elongation. This observation is supported by the reduced occupation of RNAPII over the 3’ end of selected targets. Overall, we establish MTA as a factor necessary to maintain RNAPII transcription status over a select set of genes important for proper cold stress response.

## RESULTS

### m^6^A marks do not correlate with mRNA abundance after cold exposure

Changes in the mRNA steady-state levels have been linked to m^6^A with evidence supporting both an increased or decreased mRNA abundance linked to m^6^A (12, 13, 19, 28). We reasoned that comparing active transcription (plaNET-seq) with the steady-state levels (RNA-seq) can provide important insights into the degradation kinetics of mRNAs (Figure 1A). We used this approach to determine if m^6^A had a genome-wide stabilizing or de-stabilizing effect on mRNA as suggested previously. In a recent study, cold stressed Arabidopsis plants were probed for variations in their transcriptome wide m^6^A levels (12). We used this data along with publicly available plaNET-seq data (37) and cross examined it with our own RNA-seq data from similarly grown and treated samples (Data S1). As observed by Govindan *et al.* (12) some mRNAs gained m^6^A methylation under cold stress while others lost the m^6^A mark. We used this differential m^6^A pattern and named those genes that gained methylation (m^6^A - enriched) and those that lost methylation (m^6^A-depleted) in response to cold treatment. We then compared those genes with DE genes in plaNET-seq and RNA-seq after 12h of cold exposure versus 22°C (Figure 1B-E). Our expectation was that if the presence of m^6^A had a positive correlation with mRNA abundance, we should see a large overlap of m^6^A enriched and RNA-seq up-regulated genes. Similarly, we would see an overlap between m^6^a depleted and RNA-seq down regulated genes. Further, we should observe a discrepancy between plaNET-seq and RNA-seq since RNA stability is not reflected in active transcription measurements. However, this was not the case (Figure 1B-E). By far, the largest overlap we saw, in all cases (UP/DOWN genes vs. m^6^A-enriched and UP/DOWN genes vs. m^6^A -depleted), was between plaNET-seq and RNA-seq UP/DOWN genes. This suggested that neither enrichment nor depletion of m^6^A after cold exposure correlated well with RNA abundance, an observation corroborated by recent a study (30).

**Figure 1.**
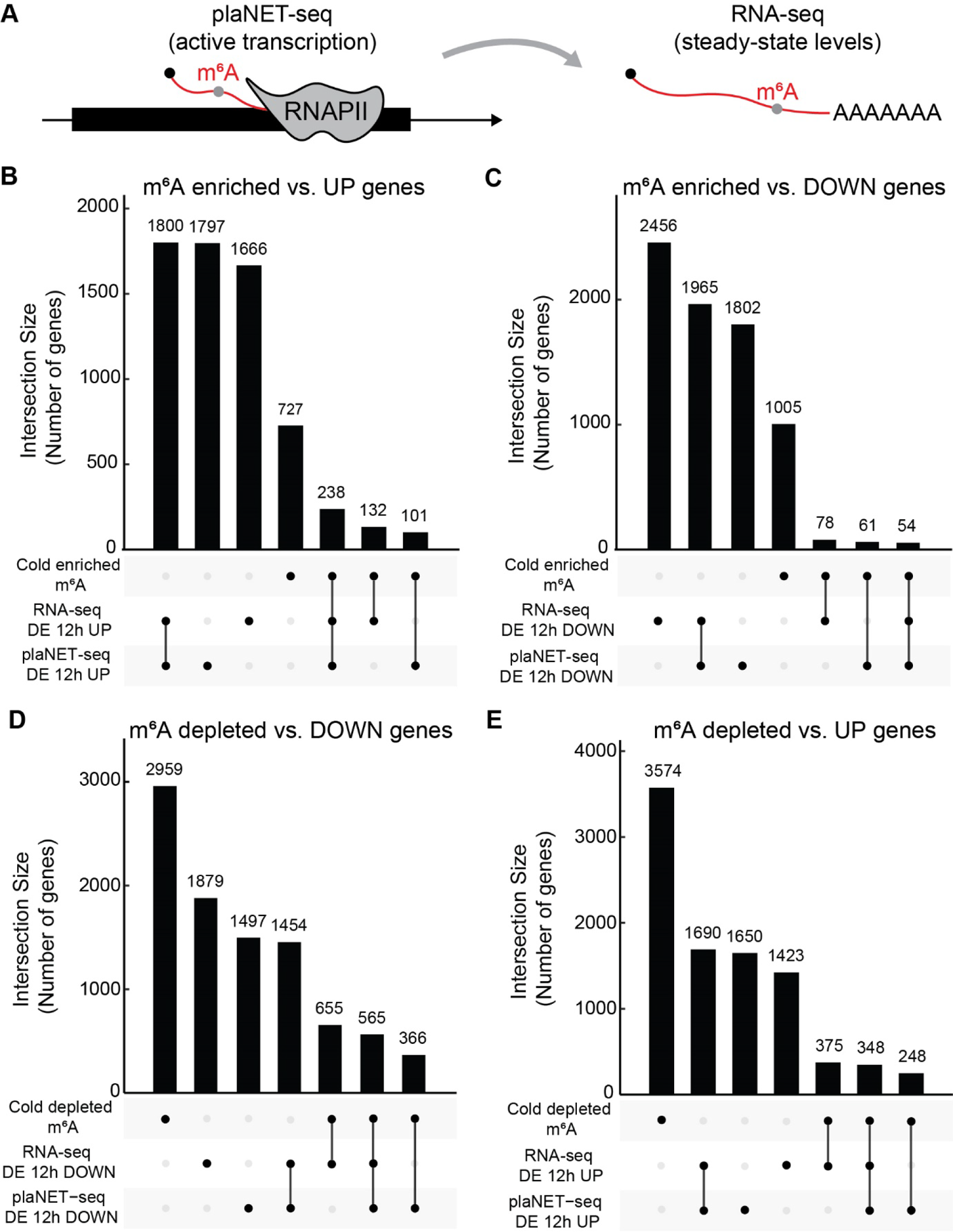
m^6^A marks do not correlate with increased RNA abundance during cold exposure. **A)** Graphical illustration of what is measured by plaNET-seq and RNA-seq. plaNET-seq measures active transcription via sequencing of RNAPII-associated RNA. RNA-seq measures the steady-state levels of RNA. **B-E)** UpSet graphs of the interaction between **B)** UP genes plaNET-seq, UP genes RNA-seq, and m^6^A enrichment, **C)** DOWN genes plaNET-seq, DOWN genes RNA-seq, and m^6^A enrichment, **D)** UP genes plaNET-seq, UP genes RNA-seq, and m^6^A depletion, and **E)** DOWN genes plaNET-seq, DOWN genes RNA-seq, and m^6^A depletion. In all comparisons, the largest interaction was between plaNET-seq and RNA-seq data.

Since m^6^A deposition is known to be co-transcriptional in animals (15, 31) and Arabidopsis MTA interacts with RNAPII (11), we inquired whether the observable differences in mRNA abundance may be a consequence of different transcription rates. Interestingly, cold-depleted m^6^A genes and cold-enriched m^6^A genes tended to be down- and up-regulated in their active transcription after cold exposure, respectively (Figure 2A-B), indicating a correlation between m^6^A and level of active transcription. To confirm a direct role in active transcription, we extracted all canonical m^6^A motifs (RRACH) with confirmed methylation (12) and compared those with motifs that do not get methylated. Intriguingly, we could detect a clear RNAPII stalling site in the plaNET-seq data for methylated sites that spanned around 100 base pairs up- and downstream of the motif (Figure 2C). Stalling was not detected around motifs that ended up non-methylated (Figure 2D). Furthermore, active transcription level was higher for those motifs that are methylated compared to those that do not (Figure 2C-D). We could not see any stalling around the other prevalent RNA methylation motif UGUA (Figure S1A), suggesting a nucleotide-specific contribution to RNAPII stalling around m^6^A motifs.

**Figure 2.**
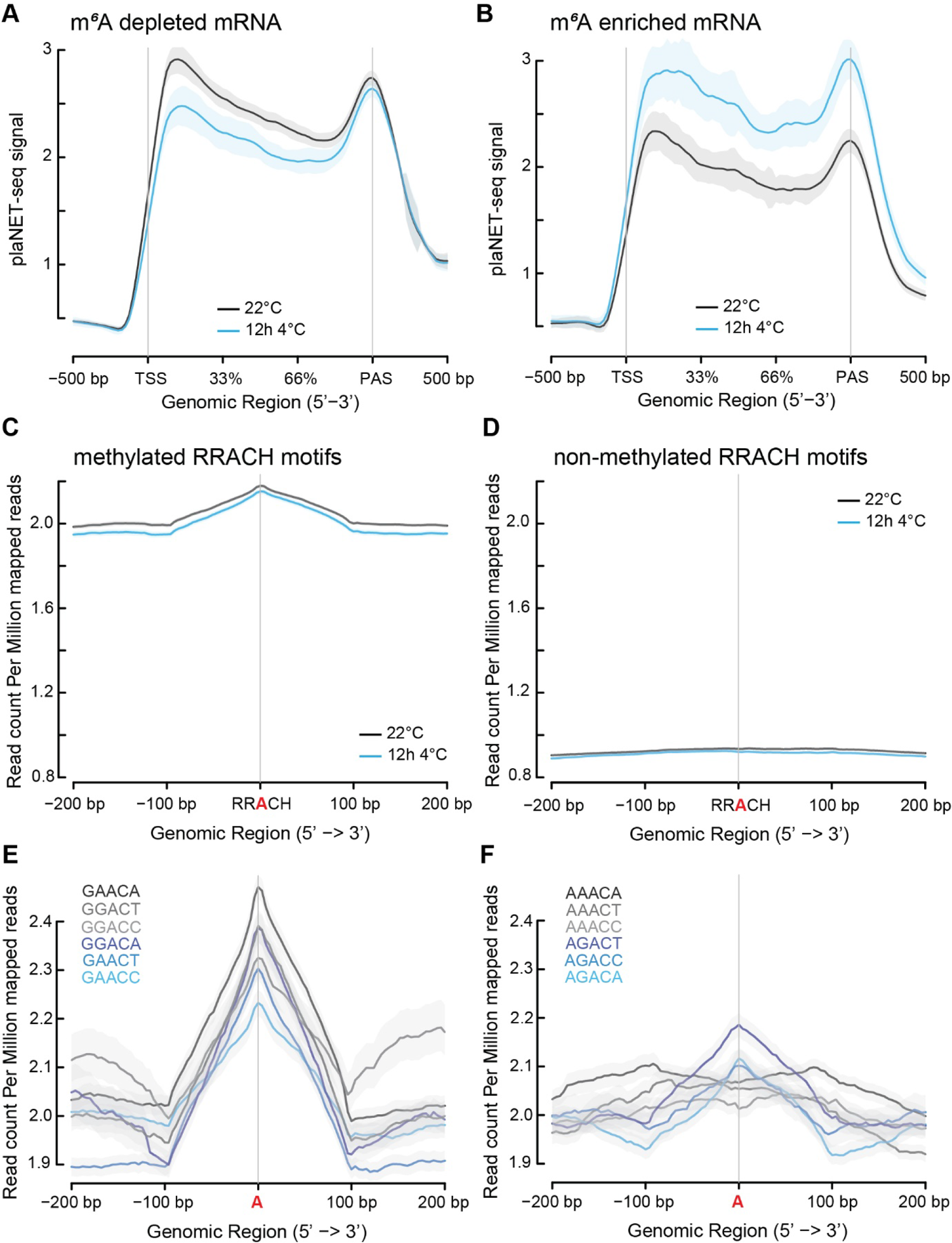
m^6^A marks correlate with transcriptional activity and stalling. ***A-B)*** Metagene profiles using plaNET-seq data of **A)** m^6^A depleted and **B)** m^6^A enriched genes at 22°C and after 12h at 4°C. Graphs show the gene body ±500 bp. Genes with m^6^A depleted mRNA tend to be down-regulated after cold exposure while genes with m^6^A enriched mRNA tend to be up-regulated after cold exposure. ***C-D)*** Metagene profiles using plaNET-seq data centered around RRACH motifs of **C)** methylated and **D)** non-methylated motifs at 22°C and after 12h at 4°C. Graphs show the RRACH motif ±200 bp. Motifs that get methylated show a clear RNAPII stalling site around the RRACH while non-methylated sites show no stalling. ***E-F)*** Metagene profiles using plaNET-seq data centered around the methylated A of **E)** 5’-G motifs and **F)** 5’-A motifs at 22°C. Graphs show the methylated A ±200 bp. 5’-GG contribute most to RNAPII stalling while 5’-AA show no stalling.

Thus, we separated each combination (12 in total for RRACH) and produced new metagene profiles. Motifs starting with a G tended to contribute more to RNAPII stalling (Figure 2E) compared to those motifs that started with an A (Figure 2F). Motifs with a 5’-AA did not show any stalling at all, indicating that there is indeed a sequence specific effect on RNAPII stalling around m^6^A motifs. Moreover, the three 5’-AA motifs showed the lowest methylation frequency (Figure S1B), suggesting that RNAPII stalling is correlated with probability of RNA methylation. Together, these results strongly point to a novel co-transcriptional role for the m^6^A writer complex that correlates with high transcriptional activity. In addition, the m^6^A writer complex may have an important role in regulating transcription in response to cold temperatures.

### The transcriptional cold response depends on the m^6^A writer MTA

As a core subunit of the writer complex, MTA is responsible for putting m^6^A marks on newly synthesized RNA. Previous studies have identified MTA as important for the cold response in Arabidopsis (12), although the mode-of-action remains unclear. To shed some light on the involvement of MTA in the early cold response, we expanded our RNA-seq study with a *mta* hypomorph mutant and an additional time point, 3h at 4°C, to increase the time resolution.

This *mta* hypomorph mutant (called *mta* hereafter) was first described in a study by Bodi *et al.* where embryo lethal T-DNA mutant of MTA was complemented with an pABI3::MTA construct to rescue the embryo lethality. (18). The ABI3 promoter is only active in the embryo and rescues the mutant from lethality. Nonetheless, we confirmed that we had down-regulation of MTA in the *mta* mutant throughout the cold exposure series in our RNA-seq data (Figure S2A). It is to be noted that the *mta* mutant is not a complete knockout, we still detect residual full-length RNA of MTA in our data, albeit to a much-reduced level. The cold sensitive phenotype of *mta* has been previously noted in (12, 30). Nevertheless, we performed electrolyte leakage assay and confirmed that the *mta* mutant had impaired cold acclimation in our conditions as well (Figure S2B). Genome-wide, the cold treatment greatly affected the transcriptional output of both WT and *mta*, and to a similar extent (Figure 3A, Data S1, S2). However, there was a large difference between WT and *mta*, among genes that were DE (Figure 3B, Data S3). In *mta* hypomorph, 419 genes were DE at 22°C, 373 DE genes after 3h at 4°C, and 1346 DE genes after 12h at 4°C vs WT. Down-regulated genes in *mta* after 12h at 4°C included several *COLD RESPONSIVE* (*COR*) genes, such as *COR27* (At5g42900), *COR28* (At4g33980), and *COR78* (At5g52310), partly explaining the *mta* mutant’s inability to cold acclimate (Figure S3, Data S3). Overall, we could see that many of these DE genes in *mta* were involved in stress response (Figure 3C). Thus, our RNA-seq experiment confirmed that MTA is an integral part of the cold response and indicates an important role for MTA in the general stress readiness of the plant.

**Figure 3.**
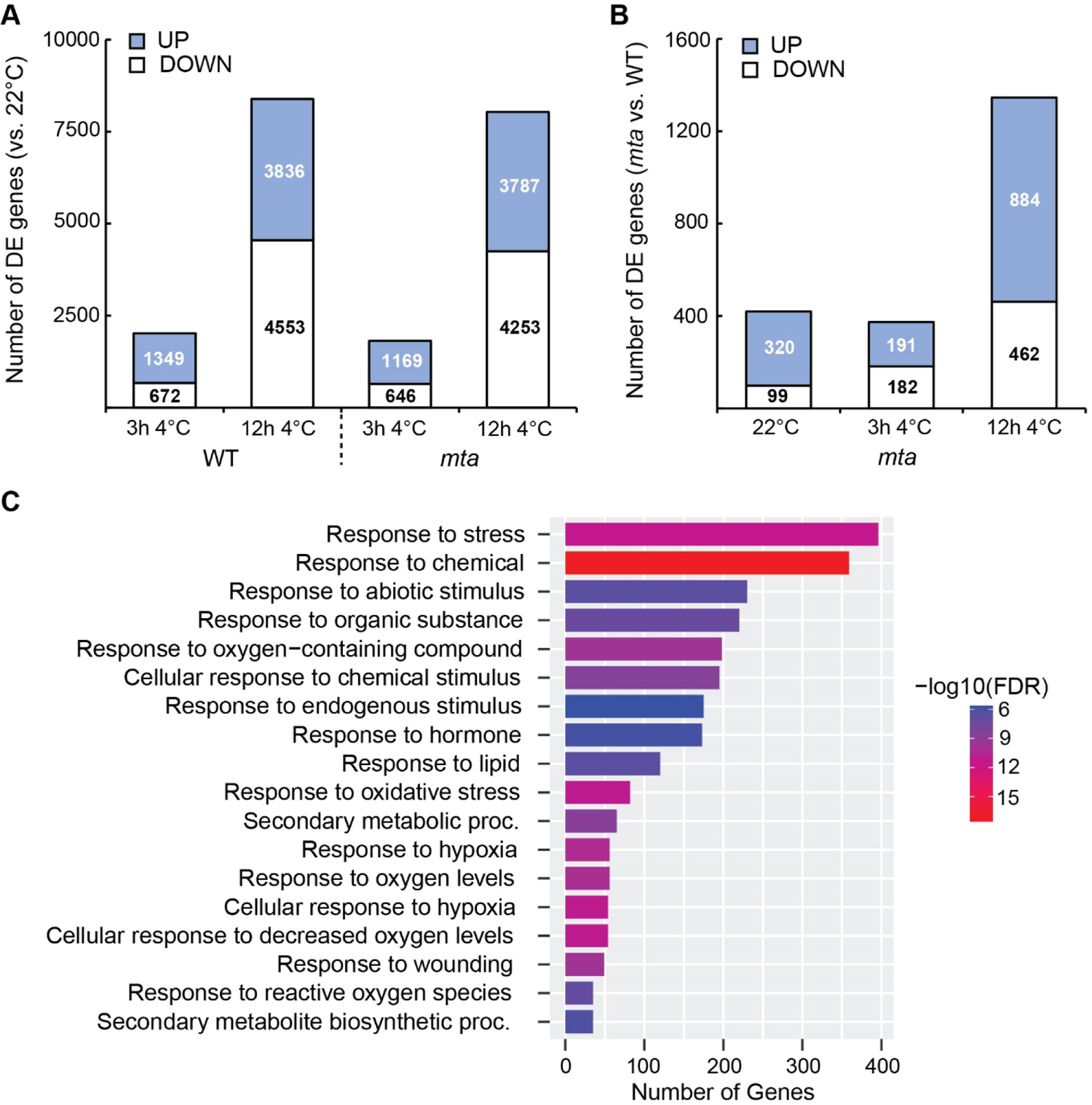
The mta mutant show impaired cold response. **A)** Number of differentially expressed genes measured by RNA-seq in wild-type and mta in response to cold stress. **B)** Number of differentially expressed genes in mta compared to wild type. **C)** GO term enrichment of differentially expressed genes in mta compared to wild type.

### MTA affects nascent transcription of the *CBFs* under cold stress

Intriguingly, when we delved deeper into the RNA-seq data, we could see that a majority of the genes that were DOWN in *mta* compared to WT at 3h and 12h were cold-induced in WT (Figure S4A-B). We detected a similar pattern when we plotted the DOWN genes in *mta* with plaNET-seq data (Figure 4A-B). DOWN genes in *mta* were greatly induced by cold temperature on a nascent transcription level in wild type. This observation was enhanced at 3h of 4°C but still present after 12h at 4°C compared to 22°C. We observed small differences of the plaNET-seq profiles for UP genes in *mta* (Figure 4C-D), indicating that MTA is primarily involved in the activation of transcription. To corroborate these results and the mis-regulation of *COR* genes to link the failed activation of cold-induced genes in *mta* with the mutant’s inability to cold acclimate, we measured the steady state levels of *CBF2* and *CBF3* at 3h 4°C. As key transcription factors in the cold response and activators of *COR* genes, *CBF2* and *CBF3* are part of the most induced genomic region early in cold exposure (27).

**Figure 4.**
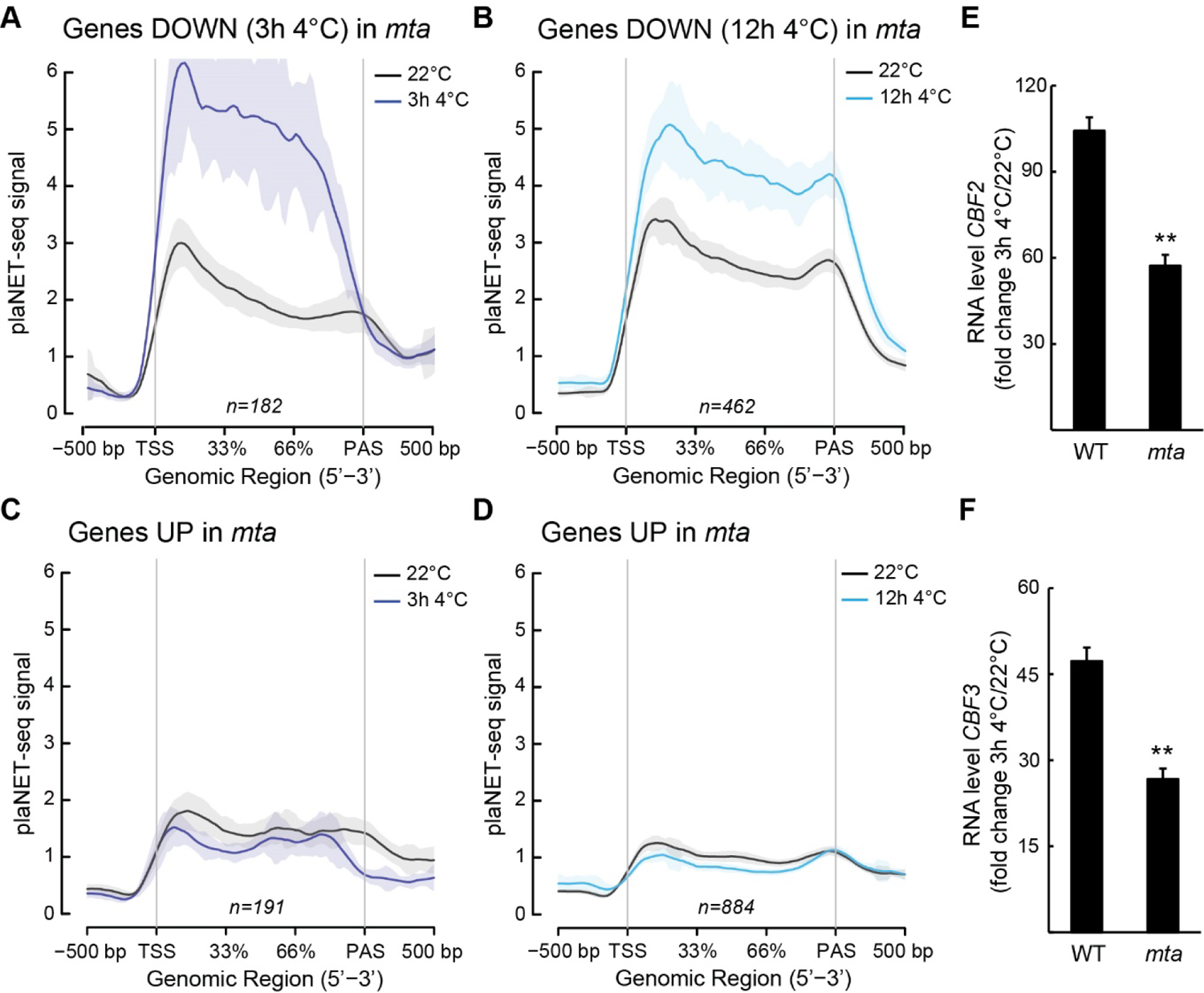
The mta mutant does not activate cold-induced genes as wild type. ***A-B)*** Metagene profiles using plaNET-seq data from wild type of genes DOWN in mta after **A)** 3h at 4°C and **B)** 12h at 4°C compared 22°C. Graphs show the gene body ±500 bp. Genes that are DOWN in mta tend to be up-regulated in wild type after cold exposure. ***C-D)*** Metagene profiles using plaNET-seq data from wild type of genes UP in mta after **A)** 3h at 4°C and **B)** 12h at 4°C compared 22°C. Graphs show the gene body ±500 bp. No difference can be seen for UP genes. ***E-F)*** The relative steady state level of **E)** CBF2 and **F)** CBF3 measured with RT-qPCR in WT and mta at 22°C compared to 3h at 4°C. Steady state levels were normalized to the WT levels at 22°C. The mean values are from three biological replicates. Error bars represent ± SEM. Statistical significance was calculated with Student’s t-test (** p<0.01).

Both genes had a significant lower induction in *mta* compared to WT after 3h of cold (Figure 4E-F). We could detect a significant decrease by qPCR, however, the level at 3h 4°C was not reduced enough to be detected by RNA-seq. These results suggest that MTA is especially important for those genes that are nascently induced by cold temperature. Both *CBF2* and *CBF3* have m^6^A marks, albeit the marks were not regulated in response to short-term cold exposure as measured by m^6^A -IP-qPCR (Figure 5A). Thus, these results confirm that MTA and the writer complex are in close proximity to RNAPII during the transcription of *CBF2* and *CBF3* at both 22°C and 4°C. Next, we aimed to measure the RNAPII occupancy to determine the active transcription of *CBF2* and *CBF3*. RNA was isolated from transcribing complexes with an RNAPII-specific antibody and used for cDNA synthesis. We used two probes along the gene bodies of *CBF2* and *CBF3*, one in each end of the genes, to see the progress of RNAPII along the genes (Figure 5B-C). The analysis revealed that at both 22°C and after 3h at 4°C, *mta* had less RNAPII engaged on *CBF2* and *CBF3* for transcription compared to wild type (Figure 5B-C). The only time point with no significant change was at the 5’-end of *CBF3* after 3h at 4°C. This suggests that MTA is required at both 22°C and 4°C for proper *CBF2* expression. The direct involvement of MTA in the methylation of *CBF2* and *CBF3* indicates that this was not a secondary effect. For *CBF3*, MTA was required for proper expression at 22°C but not at 4°. However, all RNAPII complexes did not reach the 3’-end of *CBF3* at 4°C in *mta*, which might suggest that MTA is involved in the progression of RNAPII along the *CBF3* gene body.

**Figure 5.**
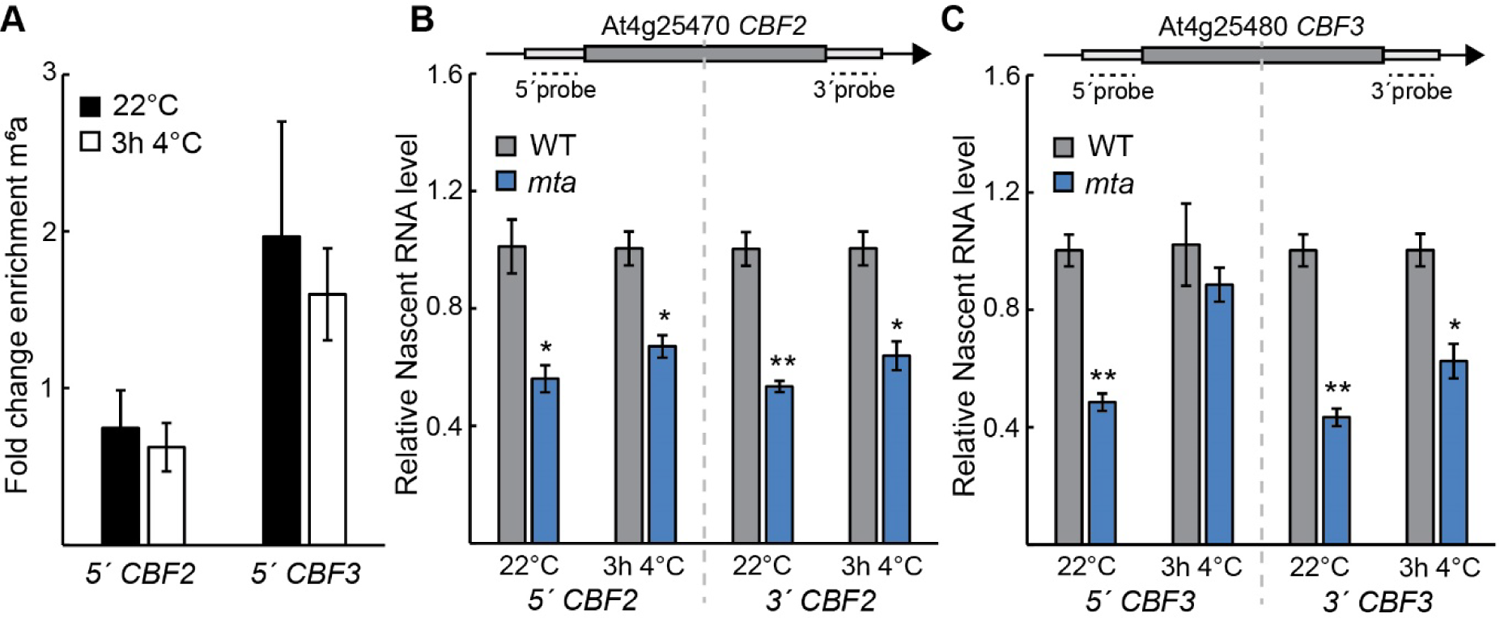
The CBFs are misregulated in the mta mutant. **A)** The level of m^6^a methylation on CBF2 and CBF3 measured by m^6^A-IP-qPCR. The mean values are from three biological replicates. Error bars represent ± SEM. No significant changes were detected between 22°C and 4°C. ***B-C)*** pNET-qPCR along the gene body of **A)** CBF2 and **B)** CBF3 at 22°C and after 3h at 4°C in wild type and mta. The mean values are from three biological replicates and normalized to the level in wild type. Error bars represent ± SEM. Statistical significance was calculated with Student’s t-test (* p<0.05, ** p<0.01).

Overall, the failed induction of the *CBFs* partly explain the *mta* mutant’s incapability to cold acclimate. Furthermore, our results from *CBF3* may indicate an additional role for MTA in the elongating RNAPII complex.

### 5’-end transcripts are accumulated for a subset of genes in the *mta* mutant

A fascinating effect of genome-wide m^6^A sites in response to cold is that the marks accumulate closer to the 5’-ends of transcripts compared to control conditions (12). Thus, we asked if the re-organizing of the m^6^A patterning in cold could influence or be a result of transcription dynamics and/or the RNA isoform composition in *mta*. This would explain the results we obtained from the *CBF3* locus. When we included plaNET-seq data from 3h at 4°C around methylated RRACH motifs, we detected a broadening of the RNAPII stalling peak, compared to 22°C and 12h at 4°C (Figure 6A). This suggests that the co-transcriptional reliance of MTA is adjusted during cold exposure at motifs that get the methylation mark.

**Figure 6.**
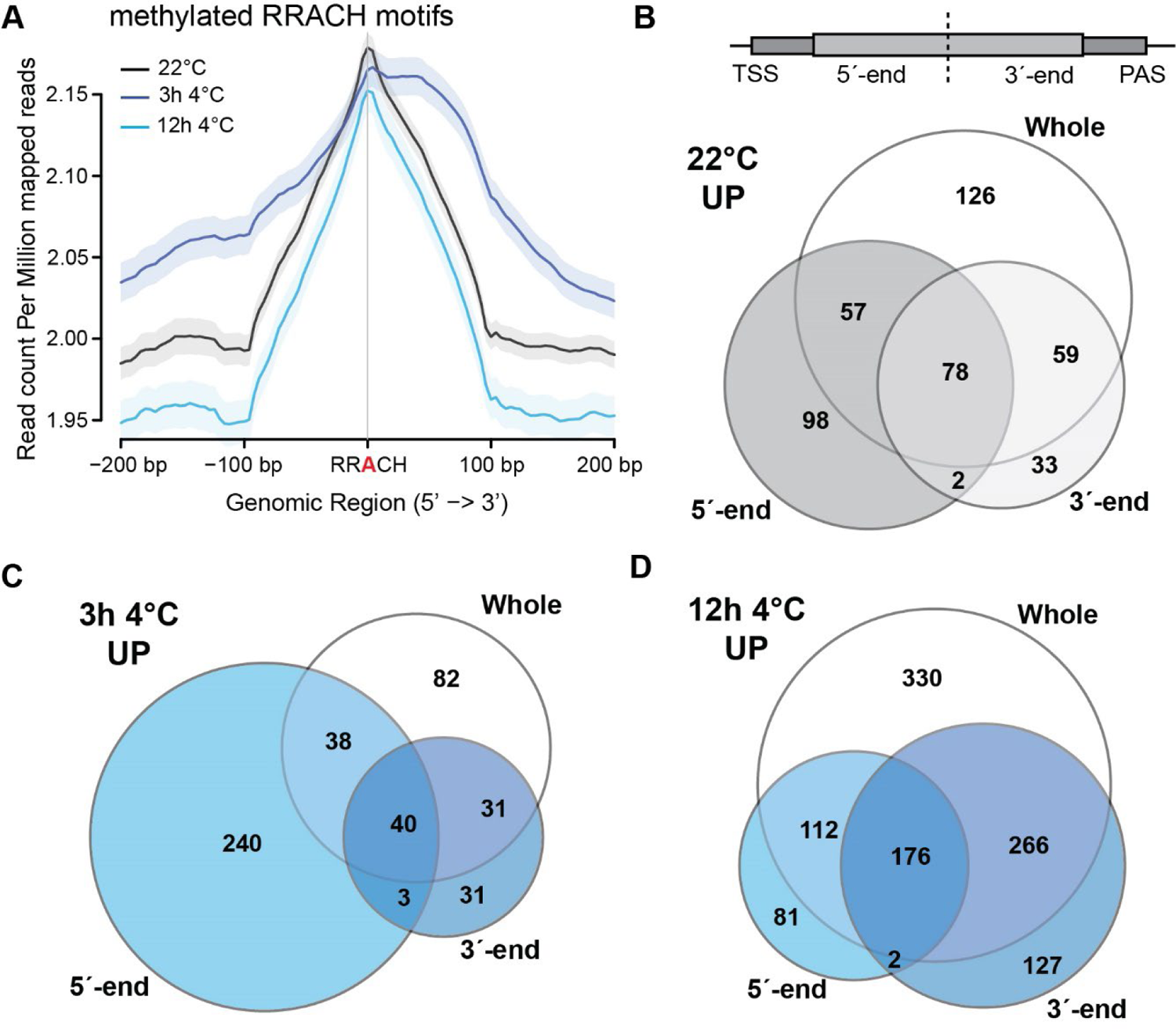
The *mta* mutant accumulates 5’-end isoforms at a subset of genes after cold exposure. **A)** Metagene profiles using plaNET-seq data centered around RRACH motifs at 22°C, 3h at 4°C and after 12h at 4°C. Graphs show the RRACH motif ±200 bp. Methylated motifs show a broader RNAPII stalling around the RRACH after 3h at 4°C compared to the other time points. ***B-D)*** Venn diagrams of differentially expressed genes (divided in the middle) between wild type and mta. Diagrams show DE genes at **B)** 22°C and after **C)** 3h at 4°C, and **D)** 12h at 4°C. There is a clear trend of more 5’-end UP genes in mta after 3h at 4°C.

Furthermore, an increase of RNAPII stalling multiplies the probability for premature termination of transcription [reviewed in (32, 33)]. If MTA is missing, it might result in accumulation of 5’-end transcripts in the *mta* mutant since co-transcriptional RNA methylation is impaired.

To test this hypothesis, we split the transcript annotation for each gene in the middle creating two transcripts for each gene i.e., 5’ end and 3’ end. Due to the resolution and 150bp insert size of our libraries we only considered transcripts over 500bp long. Then, we re-ran our RNA-seq analysis with the gene’s 5’-end and 3’-end fragments (Figure 6B, upper panel). A premature termination would show up most clearly in a 5’-end specific upregulation in *mta* compared to WT. Indeed, the number of 5’-end UP genes in *mta* showed a clear trend, they were enriched after 3h at 4°C compared to 22°C and 12h at 4°C (Figure 6B-D), correlating with the changed transcription dynamics at 3h 4°C (Figure 6A). In total, 321 genes were 5’-end UP in *mta* compared to WT (Data S4). Out of these, 240 5’-end UP genes at 3h 4°C showed no overlap with DE at whole gene or at the 3’-end. This number compared to 98 genes at 22°C and 81 genes after 12h at 4°C. In contrast, there was no clear trend of the 3’-ends on response to cold (Figure 6B-D). This supported our hypothesis that a subset of genes accumulates 5’-end isoforms in *mta* during early cold response. It further indicates that these genes amass pre-maturely terminated transcripts that do not develop into full length mRNAs.

### MTA influences elongating RNAPII at specific loci under cold stress

We reasoned that the subset of genes that accumulate 5’-end isoforms in *mta* would include good candidate genes that corroborate our hypothesis of MTA influencing RNAPII elongation. Overall, of the genes in this subset, 251 of 321 (78%) contained m^6^A marks [MeRIP-seq data from (12)](Figure 7A), indicating a direct role for MTA at many of these loci. Our final criterium for candidate genes was a cold-induced enrichment of m^6^A marks at the 5’-half of the gene, as this would ensure an increase of MTA activity on this gene end in cold. We identified 25 genes that fit our criteria (Figure 7B, Data S5). We chose 3 genes from this list for further study that showed a specific and significant increase of 5’-end fragments after 3h at 4°C in *mta* compared to WT (Figure 7C-E). BLOCK OF CELL PROLIFERATION 1 (BOP1) is a ribosome biogenesis factor (34). MERISTEM LAYER 1 (ATML1) is a homeodomain-leucine zipper class IV transcription factors involved in cell differentiation (35). Lastly, RESISTANCE TO PSEUDOMONAS SYRINGAE PV MACULICOLA INTERACTOR 1 (RIN1) has a described role in disease resistance (36). We first performed m^6^A-IP-qPCR to confirm the enrichment of methylation. We fragmented the RNA before the immunoprecipitation and used 5’-end probes to specifically detect m^6^A marks in the beginning of the genes of interest.

**Figure 7.**
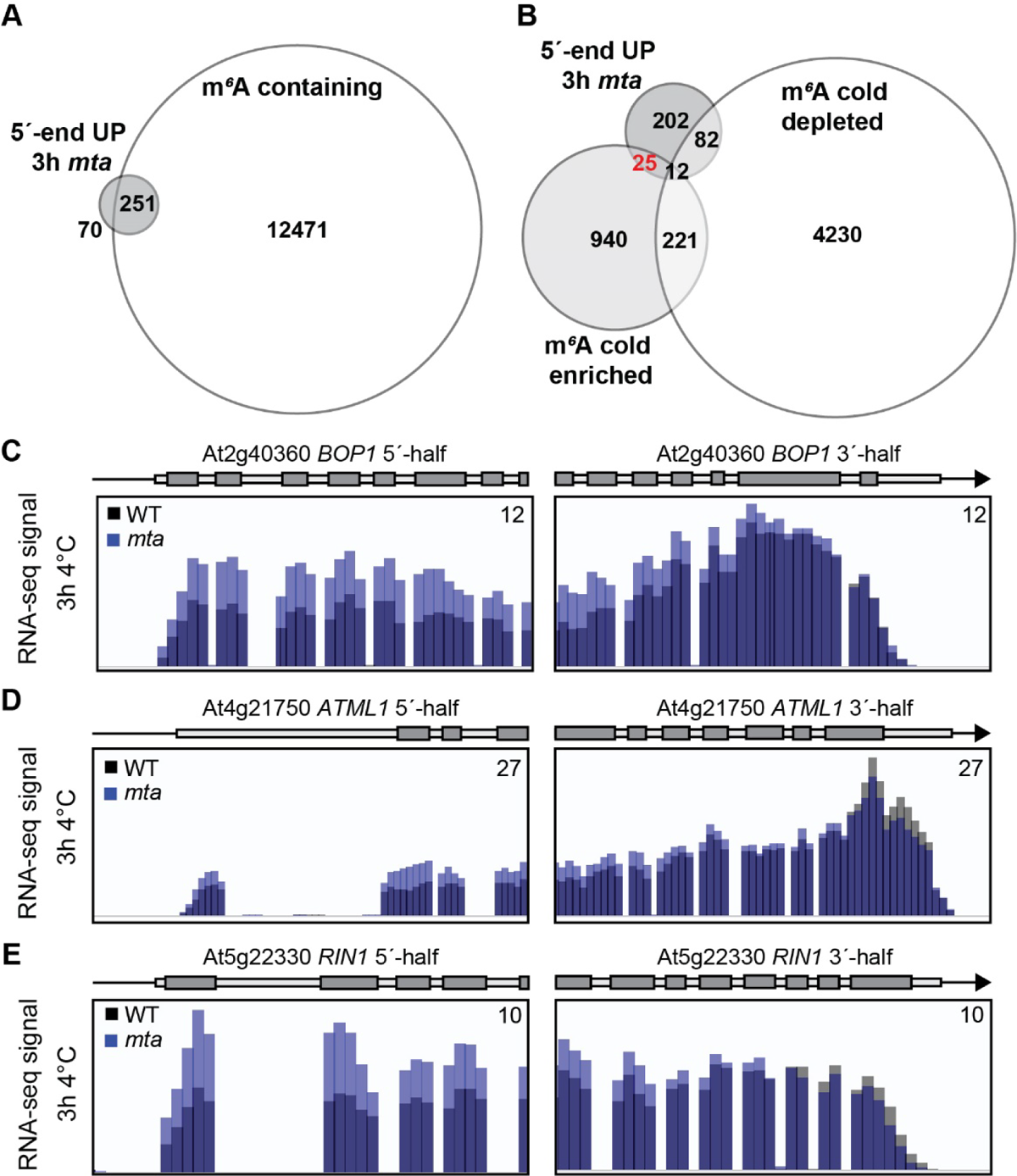
Selection of candidate genes. **A)** Venn diagram comparing 5’-end UP genes in mta with the presence of m^6^a marks. 251 out of 321 5’-end UP genes’ mRNA contain one or more marks. **B)** Venn diagram comparing 5’-end UP genes in mta with the depletion or enrichment of m^6^A marks in response to cold. 25 genes’ mRNA that are 5’-end UP in mta have an enrichment of marks in response to cold. ***C-E)*** Screenshots of RNA-seq data for candidate genes divided into 5’- and 3’-halves of the genes. RNA-seq data from three biological replicates have been merged and wild type and mta data overlayed in IGV. Screenshots show **C)** BOP1 (At2g40360) **D)** ATML1 (At4g21750), and **E)** RIN1 (At5g22330).

After 3 hours at 4°C, our candidate genes showed an enrichment of m^6^A (Figure S5A-B), indicating increased MTA activity on the 5’-end of these genes early in the cold treatment. Thus, more m^6^A marks correlated with accumulated 5’-half isoforms at our candidate genes in *mta*. We could now test the hypothesis that MTA influences RNAPII elongation at these loci by measuring the RNAPII activity on them. If the hypothesis holds true, we should detect similar levels of transcribing RNAPII in the 5’-end of the genes between WT and *mta*, but a *mta*-specific decrease of RNAPII activity at the 3’-end at 4°C compared to 22°C. If the hypothesis is false, we should detect increased RNAPII activity in *mta* compared to WT in the beginning of the genes at 4°C, explaining the increased short isoforms at the 5’-half. For our three candidate genes, we designed oligos in the 5’-end of the genes and could not detect any significant difference at the 5’-end on a nascent RNA level between WT and *mta* at 22°C or 3h at 4°C (Figure S5C-E) and a similar regulation within the genotypes (Figure 8A-C), indicating no mis-regulation on the nascent transcription level between *mta* and WT at the 5’-end of our candidate genes. In contrast, at the 3’-end, we detected a significant decrease of RNAPII activity at 4°C compared to 22°C in the *mta* mutant but not in WT (Figure 8D-F).

**Figure 8.**
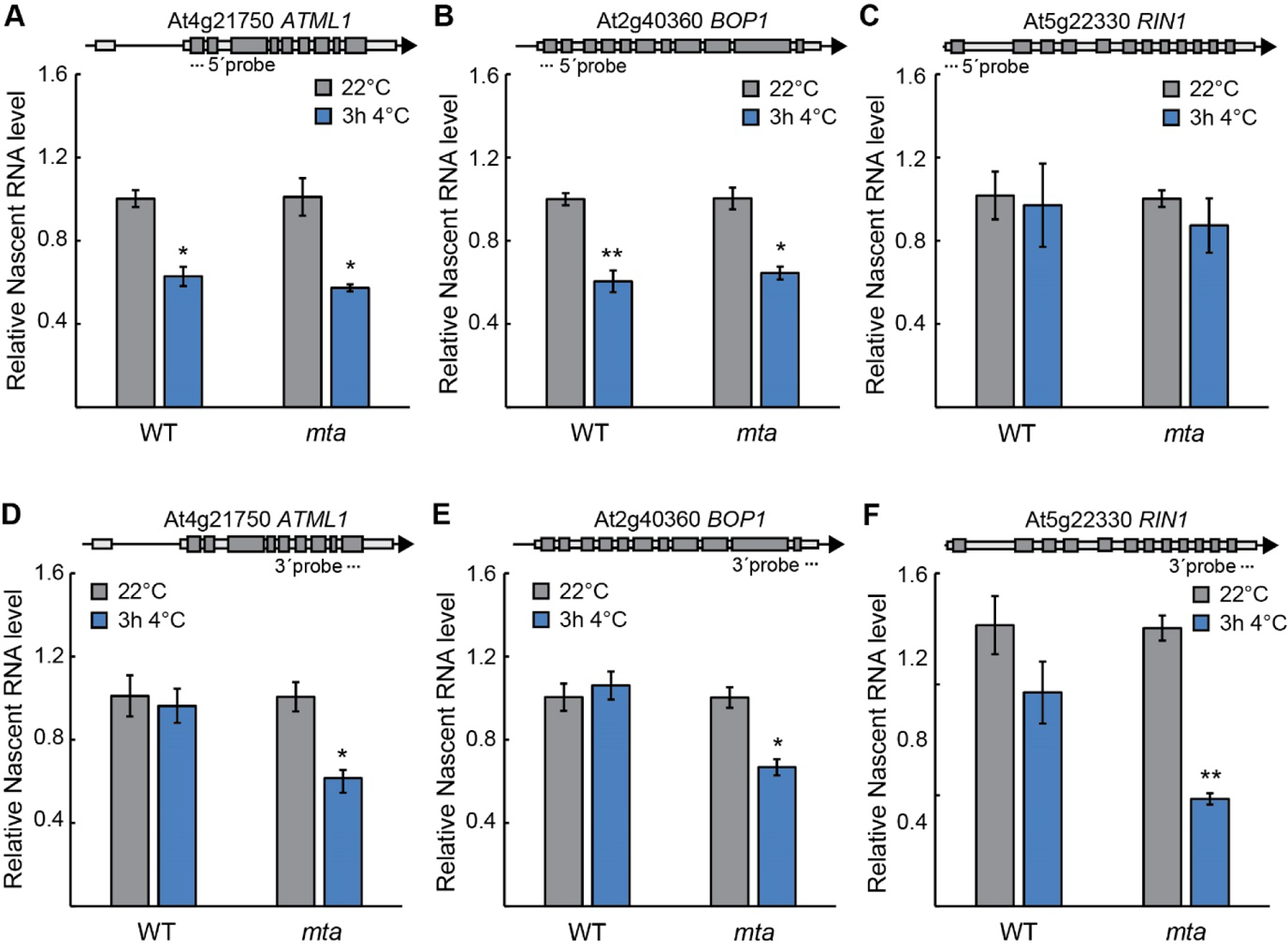
MTA influences transcription elongation for a subset of genes. ***A-C)*** pNET-qPCR of the 5’-end of, **A)** BOP1, **B)** ATML1, and **C)** RIN1 at 22°C and after 3h at 4°C in wild type and mta. The mean values are from three biological replicates and normalized to the level in wild type. Error bars represent ± SEM. Statistical significance was calculated with Student’s t-test (* p<0.05, ** p<0.01). ***D-F)*** pNET-qPCR of the 3’-end of, **D)** BOP1, **E)** ATML1, and **F)** RIN1 at 22°C and after 3h at 4°C in wild type and mta. The mean values are from three biological replicates and normalized to the level in wild type. Error bars represent ± SEM. Statistical significance was calculated with Student’s t-test (* p<0.05, ** p<0.01).

Since *mta* does not accumulate RNAPII activity at the 5’-end compared to WT, but rather showed a decreased activity at the 3’-end, we can conclude that the increase of steady state RNA levels at the 5’-end was most likely due to pre-maturely terminated RNAPII transcription. Thus, these results corroborated our hypothesis and strongly suggest that MTA influences transcription elongation in response to cold at the genes investigated. All in all, our study identifies MTA as a key co-transcriptional component in the response to cold in plants.

## DISCUSSION

MTA is known to be a protein critical for plant life. However, it remains unclear whether the necessity for MTA stems solely from its ability to deposit m^6^A on RNA or if MTA plays a role that precedes m^6^A deposition. Evidence supporting the latter are available in the form of interactions between MTA (and its animal homolog METTL3) and RNAPII (11, 31).

Using cold stress as a differentiator we analysed differential expression of mature/stable RNA (RNA seq), nascent RNA (plaNET-seq) and m^6^A methylation levels (m^6^A-IP-seq) and looked for patterns among RNAs that either gained or lost m^6^A modification upon stress. We observe dramatic changes in the transcriptome after 12h of cold stress, both at nascent (7818 DE genes) and stable transcriptome levels (8389 DE genes). Compared to that, a total of 5743 gene transcripts are differentially methylated in response to cold (12). Surprisingly, these data do not show a correlation between m^6^A and RNA abundance at nascent and/or stable RNA level. In plants, m^6^A has been linked to both increased and decreased mRNA stability. It was shown that during salt stress, transcripts needed for an appropriate stress response gain methylation are more stable thanks to change in their secondary structures and inhibition in ribonucleolytic cleavage of such mRNAs (19, 20). On the contrary, m^6^A has also been linked to reduced transcript stability. In the context of salt stress, it was shown that the loss of m^6^A stabilizes negative regulators of stress response implicating m^6^A as a negative regulator of mRNA stability (13). YTHDF proteins (m^6^A readers) have been shown to promote mRNA degradation either by exosome mediated or endoribonucleolytic cleavage (37–39) meanwhile another m6A reader insulin-like growth factor 2 mRNA-binding proteins (IGF2BPs) promote mRNA stability (40). Considering these data, it is not out of question that depending on the location of the m^6^A mark and the ‘reader’ protein that binds to it, m^6^A can have varied effects on transcript abundance globally. Indeed, Wang *et al.* (30) concluded that reduction or enrichment of m^6^A did not alter gene expression in response to cold. In addition, our inclusion of plaNET-seq data provided a unique insight and shows that transcripts that gain m^6^A methylation in the cold (cold enriched) are simply transcribed more (have higher RNAPII occupancy throughout the gene body) and those that lose methylation (cold depleted) have lower rates of transcription. This observation provides an alternative view to the observation of higher abundance of transcripts that gain methylation under stress as seen by previous studies.

Adenosine methylation occur in a specific sequence context, the RRACH motif. It is common, however curiously, not all motifs are methylated. The differentiating factor among the methylated and the non-methylated motif remains elusive. Recently, it has been shown that the patterned m^6^A methylation generally observed in animal systems can be attributed to the proximity of a given motif to a splice junction. Exon junction complex, seemingly through a member protein eukaryotic initiation factor 4a-III (EIF4A3), suppresses m^6^A deposition in its vicinity (41, 42). We observe differential RNAPII stalling over RRACH motifs that are methylated as compared to those that are not. RNAPII showed stalling only around the methylated motifs (they also had higher rates of transcription). Moreover, the stalling patterns were found to be sequence dependent with stalling being much more prominent around motifs with a 5’G as compared to 5’A containing motifs, while motifs with 5’AA showed no stalling at all. RNAPII stalling at splice sites has been well documented in plants (29, 43). This stalling is even more prominent under cold stress (29). Our data combined with the previous studies points to a complex mechanism where the rate of transcription of RNAPII (influenced either by EJC or cold or by transcription factors) has direct consequences on the methylation status of transcripts. In addition, our data points to sequence specific stalling of RNAPII over m^6^A motifs adding yet another layer of control in deposition of m^6^A on transcripts. Our data paints a picture where a transcript is scanned for motifs (most likely by RNAPII associated MTA) and RNAPII stalling over the preferred motifs results in their methylation. Evidence of co-transcriptional deposition of m^6^A includes the proximity of MTA with RNAPII during active transcription (11). This occurs early in the elongation cycle, indicating that the RNAPII associated with MTA proceed the reading of the RRACH motif by RNAPII (11). Whether the RNAPII-MTA association is direct or mediated by newly transcribed nascent RNA demands further investigation. Moreover, animal METTL3 has been shown to bind transcription start sites of active gene promoters that contain CAAT-box element (44) while no such promoter binding ability of MTA has yet been elucidated.

Our investigation of genes differentially expressed in *mta* hypomorphic mutant revealed that, a majority of these loci are stress responsive (including cold related *CBF2* and *CBF3*) according to our GO analysis. Moreover, genes that are down-regulated in *mta* after cold exposure tend to be upregulated in WT. This indicates that MTA is important for the accumulation of cold responsive mRNAs in cold temperature. Despite *mta* being cold sensitive, we did not observe dramatic differences in the transcriptome of *mta* and WT. A possible explanation for this observation is the leaky ABI3 promoter that helps rescue embryo lethality in *mta* mutant. Consequently, *mta* expression is not completely eliminated in our mutant. In other words, the effects we see in the *mta* mutant is probably an underestimation of the de facto function of MTA. Nevertheless, *CBF2* and *CBF3* transcripts are downregulated in *mta* mutant despite no observable changes in their m^6^A methylation status under cold stress. Instead, we show that this downregulation is a consequence of lower RNAPII occupancy of these genes which is observed regardless of the cold stress.

Cold induced changes in RNAPII transcription rates have a distinct pattern where RNAPII stalling around the first nucleosome is prominent after 3h at 4°C but the stalling is abolished after 12h at 4°C (29). Intriguingly, we also observed a distinct stalling pattern over the RRACH motif after 3h at 4°C. Increased stalling of RNAPII is strongly associated with increased termination of transcription (27, 33). We were able to connect elevated stalling of RNAPII around methylated motifs to overaccumulation of 5’ ends of transcripts in the *mta* mutant. These results strikingly follow the trend of increased RNAPII stalling after 3h of stress as compared to control and 12h of stress. We were able to show that RNAPII occupancy on select genes (that gain methylation under cold stress) is reduced after 3h at 4°C only at the 3’ end. RNAPII transcription rate is affected by a variety of factors and is accompanied by various co-transcriptional processes most notably splicing. Transcription elongation factors (TEFs) affect the splicing pattern either by interacting with various splicing factors (45) or by influencing RNAPII kinetics [reviewed in (46)]. Usually, a slower transcription rate is associated with exon inclusion while faster transcription rates result in exon skipping [reviewed in (47)]. In animals, m^6^A has been linked to splicing albeit with contrasting results (17, 48). In Arabidopsis, Wong *et al.* found little to no effect of m^6^A on alternative splicing (49). Our data indicates that MTA is needed for the elongation of RNAPII over a gene body. In absence of MTA, RNAPII may be able to initiate transcription however, transcription cannot proceed through whole gene body and is terminated prematurely, resulting in the loss of RNAPII occupied transcripts. Whether MTA can affect splicing via its influence on RNAPII kinetics remains to be investigated.

In summary, our study provides novel evidence establishing MTA’s role in RNAPII elongation. The recent studies linking EJC and m^6^A deposition fortify our observations that RNAPII stalling can have direct impacts on the methylation status of transcripts. Our data suggests that MTA plays a role in the aforementioned RNAPII stalling and thus affects RNA splicing, stability etc. not just by the virtue of depositing m^6^A on the target transcripts. To what extent does MTA affect splicing and stability and whether EJC is involved in such a regulation are questions open for further inquiry. Finally, our study was designed in context of genes whose transcripts are m^6^A methylated and show observable changes in their methylation status’. A considerable number of genes whose transcripts did not show any overall changes in their methylation status’ in response to cold stress have also been identified (12). In addition, there are genes whose transcripts are not m^6^A methylated at all. Whether MTA affects RNAPII kinetics over these gene bodies remains to be seen.

## MATERIAL AND METHODS

### Plant material, growth conditions and treatments

Arabidopsis thaliana wild type (Col-0) and hypomorphic *mta* mutant (18) were used in this study. Seeds were sterilized using 70% ethanol and 0.05% Triton-X, air dried and sown on ½ MS media pH 5.7 and 1.2% agar. Plates were kept in the dark at 4°C for 48h for stratification following which they were moved to a growth chamber with ̴100µEm-2s-1 light on long day (16h light/8h dark, 22⁰C Day/18⁰C night) conditions. 12-day old seedlings were collected either directly or post cold treatment (4°C with ∼20-25µEm-2s-1 light). Three independent biological replicates were collected, flash frozen in liquid nitrogen and stored at −80°C till further use.

### RNA isolation, cDNA preparation and qPCR

RNA was isolated from ground tissue using RNeasy Plant Mini Kit (QIAGEN) according to manufacturer’s instructions. RNA thus obtained was treated with TURBO® DNAse (Thermo Fischer) according to the standard protocol. The quality of RNA was assessed using traditional agarose electrophoresis as well as Agilent Bioanalyzer 2100 system (for sequencing samples). For cDNA synthesis iScript™ cDNA Synthesis Kit (Bio-Rad) was used. cDNA was synthesized using 500ng of RNA as per manufacturer’s protocol. qPCR was performed using iTaq Universal SYBR Green Supermix (Bio-Rad) and Bio-Rad CFX96/CFX384 Touch Real-Time PCR Detection Systems. For differential expression analysis ΔCt was calculated (Ct of target – Ct of reference) and ΔΔCt was calculated (ΔCt mutant – ΔCt WT) followed by fold change (2^-ΔΔCt). p-values were calculated using Student’s t-test. All oligos used in this study can be found in Table S1.

### RNA sequencing and analysis

High quality RNA (RIN > 8) was sent to Novogene Europe for library preparation and sequencing. Strand specific libraries were generated at Novogene after polyA enrichment. Sequencing was performed on Illumina’s NovaSeq 6000 platform and 6GB of raw data was obtained per sample. Salmon files were generated using the pipeline developed at UPSC and documented here (http://franklin.upsc.se:3000/materials/materials/Guidelines-for-RNA-Seq-data-analysis.pdf). Briefly, data was pre-processed and quality checked FastQC v0.11.9 (quality control of the raw data) and SortMeRNA v4.3.4 (filter and remove rRNA contamination) (50, 51). Thereafter, Trimmomatic (52) was used to trim the adapter sequences and FastQC was performed again to ensure data integrity. Salmon v1.6.0 (53) was used to determine the read counts with AtRTD2 (54) as a reference. The salmon files thus obtained were used for the differential expression analysis using the 3D RNA-seq App (run locally, but also available online at https://3drnaseq.hutton.ac.uk/app_direct/3DRNAseq/) (55). Venn diagrams, intersect plots and GO analysis presented were generated using BioVenn (56), Intervene tool (57) and ShinyGO (58) with cosmetic changes.

### m^6^A-RNA IP

m^6^A IP was performed following the protocol described in (11) with few changes. polyA RNA was isolated using Dynabeads™ mRNA Purification Kit (Thermo Fisher) according to manufacturer’s protocol. 2µg of polyA enriched RNA was fragmented using NEBNext® Magnesium RNA Fragmentation Module (New England Biolabs) with incubation at 94°C for 1 min. RNA was purified after fragmentation and used for IP. IP protocol described in (11) was followed thereafter and Epimark® N^6^-Methyladenosine Enrichment Kit (New England Biolabs) was used. RNA from input and IP samples was used to prepare cDNA and qPCR analysis was performed as described above. The negative spike in control was used to normalize Ct values and ΔΔCt was calculated as ΔCt IP – ΔCt Input. p-values were calculated using Student’s t-test. Oligos are found in Table S1.

### Remapping of existing datasets

A few existing data sets were remapped and/or used in this study. They include plaNET-seq (GSE131733) and m^6^A-IP (GSE184056) (12, 29). Differentially expressed genes in plaNET-seq data were calculated as described (29).

### Metagene plots

The dataset (SRR9117170-SRR9117181) for plaNET-seq (GSE131733) was downloaded from NCBI’s Sequence Read Archive database. Raw reads were processed and mapped via STAR 2.7.10a based on the script 01-Alignment_plaNET-Seq.sh available at https://github.com/Maxim-Ivanov/Kindgren_et_al_2019 (37). The generated bam from replicates were merged by samtools merge function. RRACH motif coordinates were extracted from *Arabidopsis thaliana* TAIR10 genome using “motif search” function in CLC Main Workbench software. The extracted motif coordinates were annotated with known gene coordinates from Arabidopsis_thaliana.TAIR10.55.gff3 file downloaded from Plant Ensemble database (Data S6). The metagene plots were generated using ngs.plot.r (59) software using build-in Tair10 genome and bed files based on Data S6 and Data S1-3.

### plaNET-qPCR

Nuclei was isolated from around 3 grams of 12-day old seedlings with Honda buffer (0.44 M Sucrose, 1.25% Ficoll, 2.5% Dextran T40, 20 mM Tris –HCl pH 7.4, 10 mM MgCl2, 0.5% Triton-X, Prot. inhibitor tablet, RNase inhibitor, 5 mM DTT). The nuclear lysis and RNAPII-IP were done according to (29) with small modifications. Briefly, after lysis and DNAse I treatment, the supernatant was mixed with protein G magnetic beads (Thermo Scientific) coupled to an endogenous RNAPII antibody (8WG16, Sigma Aldrich) for 2h in 4°C. The beads were washed 4 times with wash buffer (0.3 M NaCl, 20 mM Tris-HCl pH 7.5, 5 mM MgCl2, 5 mM DTT, proteinase inhibitor tablet and RNase inhibitor (20 U/ml)). To disrupt the RNAPII complexes, QIAzol was added, and RNA was isolated using the miRNeasy kit from Qiagen. RNA concentration was measured with Nanodrop and approximately 100 ng was used for cDNA synthesis with gene specific primers and Superscript IV (Invitrogen) according to manufacturer’s instructions. Oligos used can be found in Table S1.

### Freezing test

Electrolyte leakage measurements were carried out according to (60). In short, plants were grown in short days (8h light /16 h dark cycle) for 4 weeks. For the cold acclimation experiments, WT and mutant plants were transferred to a cold chamber set at 4 ⁰C for 4 days. The freezing bath (FP51, Julabo) was set to −2⁰ C when the experiment started. After 45 minutes, icing was induced manually in each tube with a metallic stick. Temperature decrease occurred at the rate of −1 °C per 30 mins, and samples were taken out at designated temperature point(s). Later, 1.3 ml of water was added to each tube and placed on a shaker overnight at 4⁰ C and conductivity was measured using a conductivity cell (CDM210, Radiometer) on the next day. All tubes were then subjected to flash freeze and left on a shaker overnight at room temperature and measured for conductivity again. Data were fitted into a sigmoidal dose-response curve using GraphPad Prism software. Statistical significance between LogEC50 was calculated with an extra sum-of-squares F-test included in GraphPad Prism.

## ACKNOWLEDGEMENTS

We would like to thank members of the Kindgren lab for their critical reading of the manuscript. A special thanks to the Nicolas Delhomme and the bioinformatic facility at Umeå Plant Science Centre for assistance in the bioinformatic analysis and the greenhouse personnel at Umeå Plant Science Centre for plant maintenance.

## FUNDING

This work was supported by the Swedish Research Council [2018-03926 to P.K.]; and grants from the Knut and Alice Wallenberg Foundation and the Swedish Governmental Agency for Innovation Systems [KAW 2016.0355 and 2020.0240, VINNOVA 2016-00504]. Funding for open access charge: Swedish University of Agricultural Sciences.

## AUTHOR CONTRIBUTIONS

Conceptualization: SSB, PK

Methodology: SSB, DB, PK

Investigation: SSB, DB, PK

Visualization: SSB, DB, PK

Supervision: PK

Writing - original draft: SSB, PK

Writing - review & editing: SSB, DB, PK

## COMPETING INTEREST

The authors declare no conflict of interest.

## DATA AND MATERIALS AVAILABILITY

RNA Seq data produced during this study are submitted and available online at NCBI GEO under accession number GSE226622.

## SUPPLEMENTARY MATERIALS

Figures S1-5

Tables S1

Data S1-6

## Notes

### Competing Interest Statement

The authors have declared no competing interest.

