## Supplementary material for "MTA influences RNA Polymerase II transcription dynamics and regulates the cold response in Arabidopsis": Supp Figures

Susheel Sagar Bhat *et al.*

**This PDF file includes:**

Figs. S1 to S5
Tables S1

**Other Supplementary Materials for this manuscript include the following:**

Data S1 to S6

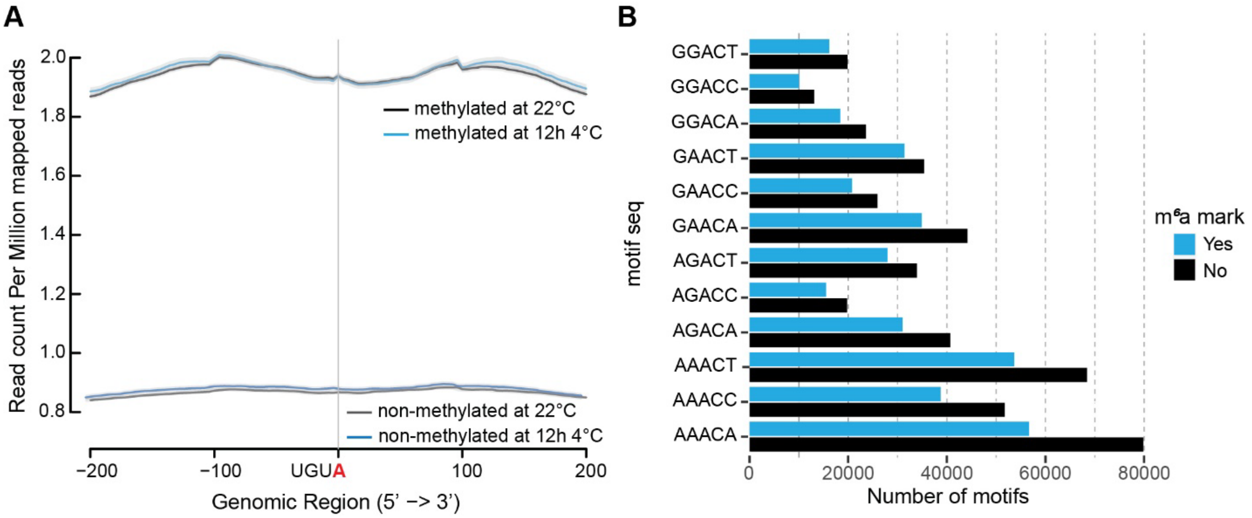


**Fig. S1.**

**A)** Metagene profiles using plaNET-seq data centered around UGUA motifs of methylated and non-methylated motifs at 22°C and after 12h at 4°C. Graphs show the UGUA motif ±200 bp.

**B)** Frequency of methylation of each combination of the RRACH motif.

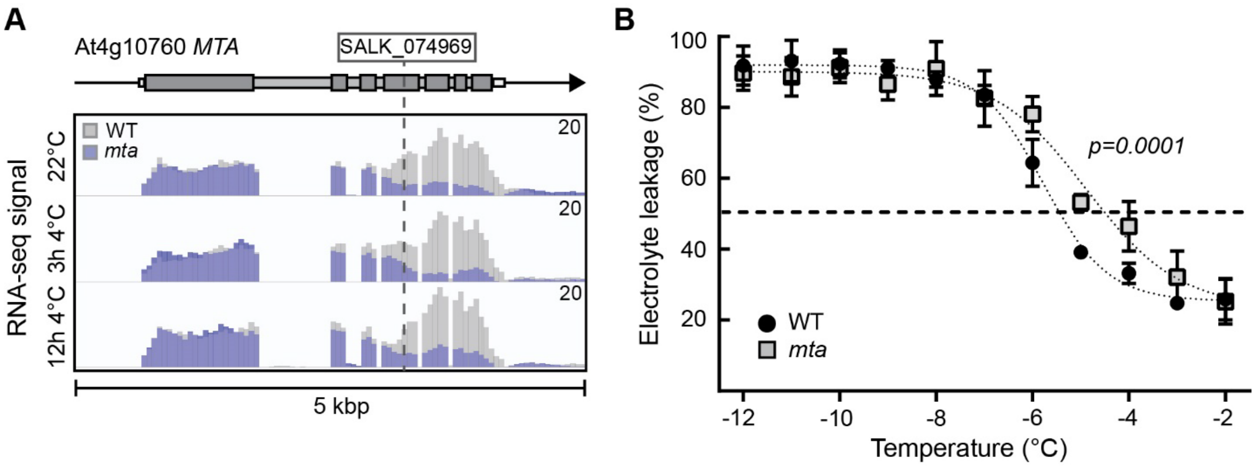


**Fig. S2.**

**A)** Screenshot of RNA-seq data for *MTA* (At4g10760). RNA-seq data from three biological replicates have been merged and wild type and *mta* data overlayed in IGV.

**B)** Electrolyte leakage in wild-type and *mta* of cold-acclimated (5 days of 4°C) plants. Each data point represents the mean from at least 3 biological replicates (±SEM).

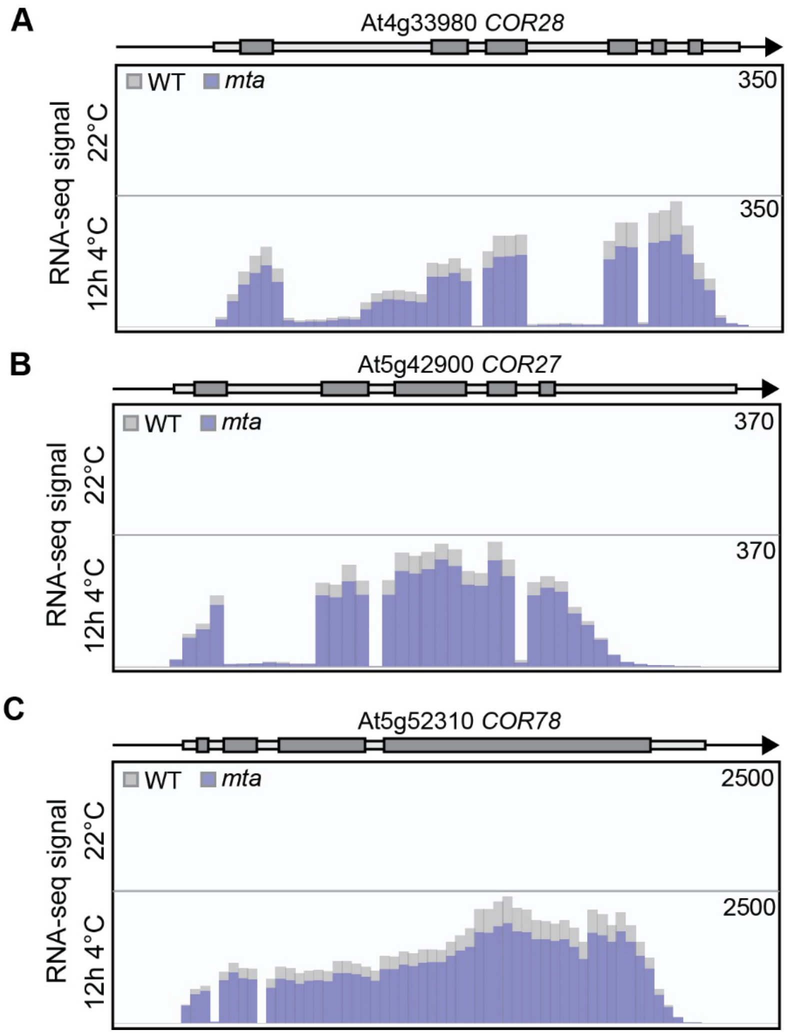


**Fig. S3.**

**A-C)** Screenshots of RNA-seq data for *COR* genes. RNA-seq data from three biological replicates have been merged and wild type and mta data overlayed in IGV. Screenshots show **A)** *COR28* (At4g33980) **B)** *COR27* (At5g42900) **C)** *COR78* (At5g52310).

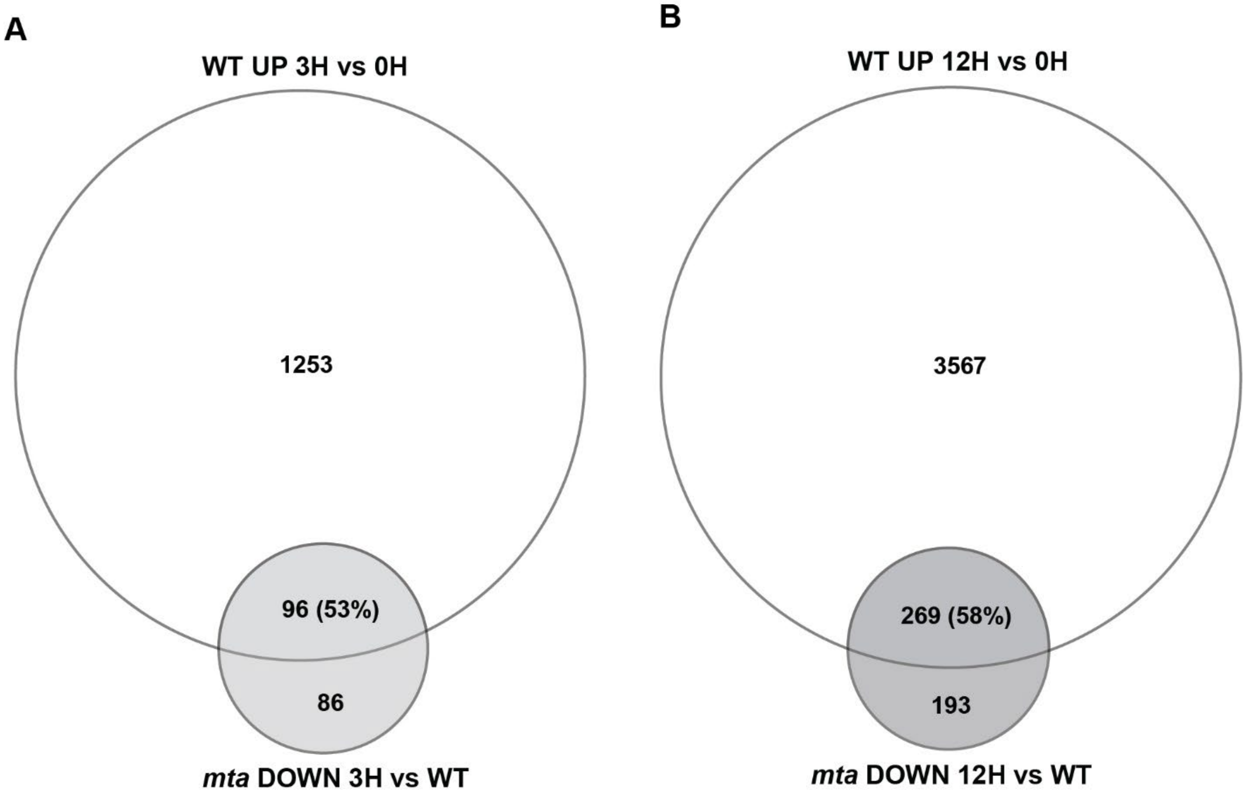


**Fig. S4.**

**A)** Venn diagram comparing UP genes in wild type with DOWN genes in *mta* after 3h at 4°C.

**B)** Venn diagram comparing UP genes in wild type with DOWN genes in *mta* after 12h at 4°C.

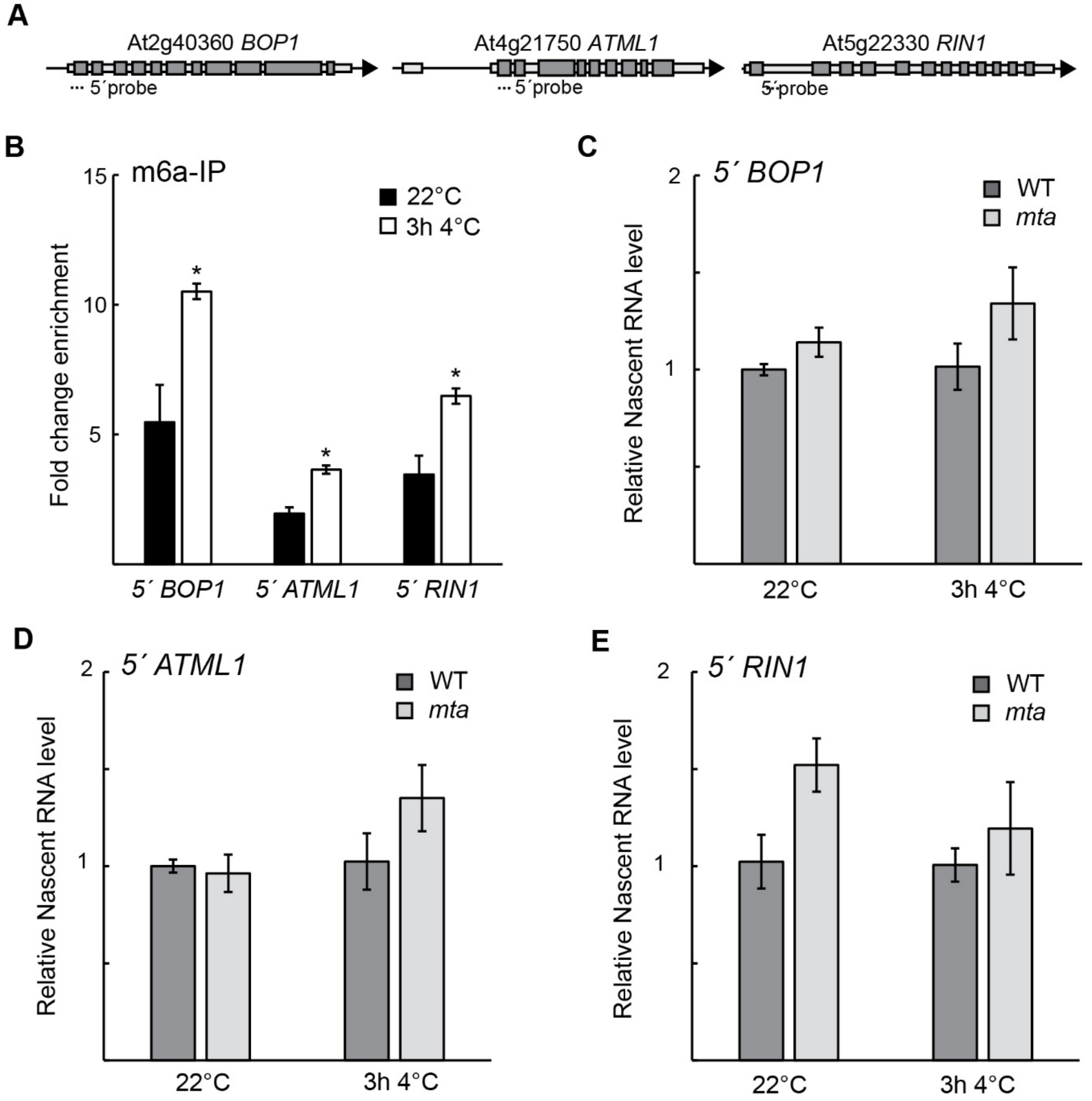


**Fig. S5.**

**A)** The level of $m^6A$ methylation on *BOP1, ATML1,* and *RIN1* measured by $m^6A$-IP-qPCR. The mean values are from three biological replicates. Error bars represent ± SEM. Statistical significance was calculated with Student's t-test (* $p<0.05$).

**B-D)** pNET-qPCR of the 5´-end of, **B)** *BOP1*, **C)** *ATML1*, and **D)** *RIN1* at 22°C and after 3h at 4°C in wild type compared to *mta*. The mean values are from three biological replicates and normalized to the level in wild type. Error bars represent ± SEM. Statistical significance was calculated with Student's t-test (* $p<0.05$, ** $p<0.01$).

**Table S1.**

A list of primers used for qPCR is provided below

| Primer Name | Primer sequence |
|---|---|
| 5'UTR CBF2 F | ATAACTTCAAACACTTACCTGAATTAG |
| 5'UTR RT CBF2 | GACTGGAGCACGAGGACACTGGTATGATAAAATATTTATTTCTCTATC |
| 3'UTR CBF2 F | TCGATTTTTATTTCCATTTTTGG |
| 3'UTR RT CBF2 | GACTGGAGCACGAGGACACTTTACATTCGTTTCTCACAACCA |
| 5'UTR CBF3 F | CAAACATTAAATCCACCTGAACT |
| 5'UTR RT CBF3 | GACTGGAGCACGAGGACACTTTTGCTGAAATAATAGTTTCTCTCTCT |
| 3'UTR CBF3 F | TTTAGAATGGAATCTTCATTATGTTTG |
| 3'UTR RT CBF3 | GACTGGAGCACGAGGACACCCCACACTTATACTGAAACTGAATC |
| ATML1 5'end_F | TCCAAACATGTTCGAATCTCA |
| ATML1 5'end_R | ACTTCTGCGCCGGACTTAG |
| ATML1 3'end_F | CGGTTCACTACTCACAGTTGC |
| BOP1_5' end_F | GGAGCAAAGGAGCTAACGAA |
| BOP1_5' end_R | AAGCTTTCTGCTTCTACTGGTTTC |
| BOP1_3'end_F | TTGTTGTATTCACCACCTTTACTCA |
| BOP1_3'end_R | CTTAACCAACGCTGCCAAAT |
| RIN1_5' end_F | CAGTCCACCGCTAAGAAACA |
| RIN1_5' end_R | CAAATCCAGCTGCCAATTTT |
| RIN1_3' end_F | TGTGCAGCTTCTGTCTCCTG |
| RIN1_3' end_R | TGCAAAAGCTTTGCTGAAGA |
| Neg_m6A_F | TCTCTGGCCTCTGTGGAGAT |
| Neg_m6A_R | AACCGTCAAACTCCTGGTTG |
| Pos_m6A_F | AGCAGTTCATCGCACAGGTC |
| Pos_m6A_R | AGCCACTTCTTGAGCAGGTC |
| rtUBQ10 F | GGCCTTGTATAATCCCTGATGAATAAG |
| rtUBQ10 R | AAAGAGATAACAGGAACGGAAACATAGT |
| rtACT2 F | CTTGCACCAAGCAGCATGAA |
| rtACT2 R | CCGATCCAGACACTGTACTTCCTT |

**Data S1. (separate file)**

A list of differentially expressed genes in Col-0 after 12H and 3H of cold stress (4°C) vs control.

**Data S2. (separate file)**
A list of differentially expressed genes in *mta* after cold stress.

**Data S3. (separate file)**
A list of differentially expressed genes *mta* vs Col-0 in control and after cold stress.

**Data S4. (separate file)**
A list of genes differentially expressed only at the 5' end in *mta* vs Col-0 after 3H of cold stress.

**Data S5. (separate file)**
25 genes that are UP regulated in mta (vs Col-0) only at the 5' end after 3H of cold stress and also gain $m^6A$ mark upon cold stress (meRIP data from Govindan et al., 2022).

**Data S6. (separate file)**
Extracted RRACH motifs along with their genomic coordinates and whether they are methylated or not.
